# What you see depends on what you hear: temporal averaging and crossmodal integration

**DOI:** 10.1101/362665

**Authors:** Lihan Chen, Xiaolin Zhou, Hermann J. Müller, Zhuanghua Shi

## Abstract

In our multisensory world, we often rely more on auditory information than on visual input for temporal processing. One typical demonstration of this is that the rate of auditory flutter assimilates the rate of concurrent visual flicker. To date, however, this auditory dominance effect has largely been studied using regular auditory rhythms. It thus remains unclear whether irregular rhythms would have a similar impact on visual temporal processing; what information is extracted from the auditory sequence that comes to influence visual timing; and how the auditory and visual temporal rates are integrated together in quantitative terms. We investigated these questions by assessing, and modeling, the influence of a task-irrelevant auditory sequence on the type of ‘Ternus apparent motion’: group motion versus element motion. The type of motion seen critically depends on the time interval between the two Ternus display frames. We found that an irrelevant auditory sequence preceding the Ternus display modulates the visual interval, making observers perceive either more group motion or more element motion. This biasing effect manifests whether the auditory sequence is regular or irregular, and it is based on a summary statistic extracted from the sequential intervals: their geometric mean. However, the audiovisual interaction depends on the discrepancy between the mean auditory and visual intervals: if it becomes too large, no interaction occurs – which can be quantitatively described by a partial Bayesian integration model. Overall, our findings reveal a crossmodal perceptual averaging principle that may underlie complex audiovisual interactions in many everyday dynamic situations.

**Public Significance Statement:** The present study shows that auditory rhythms, regardless of their regularity, can influence the way in which the visual system times (subsequently presented) events, thereby altering dynamic visual (motion) perception. This audiovisual temporal interaction is based on a summary statistic derived from the auditory sequence: the geometric mean interval, which is then combined with the visual interval in a process of partial Bayesian integration (where integration is unlikely to occur if the discrepancy between the auditory and visual intervals is too large). We propose that this crossmodal perceptual averaging principle underlies complex audiovisual interactions in many everyday dynamic perception scenarios.

**Author Note:** This study was supported by grants from the Natural Science Foundation of China (31200760, 61621136008, 61527804), German DFG project SH166 3/1 and “projektbezogener Wissenschaftleraustausch” (proWA). The data, and the source code of statistical analysis and modeling are available at https://github.com/msenselab/temporal_averaging. Part of the study has been presented as a talk in 17th International Multisensory Research Forum (IMRF, June 2016, Suzhou, China).

---

Most stimuli and events in our everyday environments are multisensory. It is thus no surprise that our brain often combines a heard sound with a seen stimulus source, even if they are in conflict. One typical such phenomenon, in a performance we enjoy, is the *ventriloquism effect* (Chen & Vroomen, 2013; Occelli, Bruns, Zampini, & Roder, 2012; Recanzone, 2009; Slutsky & Recanzone, 2001): we perceive the ventriloquist’s voice as coming from the mouth of a dummy as if it was the dummy that is speaking. Of note in the present context, audiovisual integration has not only been demonstrated in spatial localization, but also in the temporal domain. In contrast to the dominance of vision in audiovisual spatial perception, audition dominates temporal processing, such as in rhythms and intervals. As an example, think of how we tend to ‘auditorize’ a conductor’s arm movements coordinating a musical passage, or Morse code flashes emanating from a naval ship. In fact, neuroscience evidence has revealed that information for time estimation is encoded in the primary auditory cortex for both visual and auditory events (Kanai, Lloyd, Bueti, & Walsh, 2011). This is consistent with the proposal that the perceptual system automatically abstracts temporal structure from rhythmic visual sequences and represents this structure using an auditory code (Guttman, Gilroy, & Blake, 2005).

Another compelling demonstration of how auditory rhythm influences visual tempo is known as the *auditory driving effect* (Boltz, 2017; Gebhard & Mowbray, 1959; Knox, 1945; Shipley, 1964): the phenomenon that variations in auditory flutter rate may noticeably influence the rate of perceived visual flicker. This influence, though, is dependent on the disparity between the auditory and visual rates (Recanzone, 2003). Quantitatively, this influence has been described by a Bayesian model of audiovisual integration (Roach, Heron, & McGraw, 2006), which assumes that the brain takes into account prior knowledge about the discrepancy between the auditory and visual rates in determining the degree of audiovisual integration. Auditory driving is a robust effect that generalizes across different types of tasks, including temporal adjustment and production (Myers, Cotton, & Hilp, 1981) and perceptual discrimination (Welch, DuttonHurt, & Warren, 1986), and it may even be seen in the effect of one single auditory interval on a subsequent visual interval (Burr, Della Rocca, & Morrone, 2013).

It should be noted, however, that auditory driving has primarily been investigated using regular rhythms, the implicit assumption being that the mean auditory rate influences the mean visual rate. On the other hand, studies on *ensemble coding* (Alvarez, 2011; Ariely, 2001) suggest that perceptual averaging can be rapidly accomplished even from a set of variant objects or events; for example, we can quickly estimate the average size of apples in a supermarket display, or the average tempo of a piece of music. With regard to the present context, audiovisual integration, it remains an open question how the average tempo in audition quantitatively influences the temporal processing of visual events – an issue that becomes prominent as the mechanisms underlying perceptual averaging processes themselves are still a matter of debate. There is evidence that the mental scales underlying the representation of magnitudes (e.g., visual numerosity and temporal durations) are non-linear rather than linear (Allan & Gibbon, 1991; Dehaene, Izard, Spelke, & Pica, 2008; Nieder & Miller, 2003). It has also been reported that, in temporal bisection (i.e., comparing one interval to two reference intervals), the subjective mid-point between one short and one long reference duration is closer to their geometric, rather than their arithmetic, mean (Allan & Gibbon, 1991). However, it remains to be established whether temporal rate averaging obeys the principle of the arithmetic mean or the geometric mean, which might have implications for a broad range of mechanisms coding ‘magnitude’ in perception (Walsh, 2003).

On these grounds, the aim of the present study was to quantify temporal rate averaging in a crossmodal, audiovisual scenario using irregular auditory sequences. To this end, we adopted and extended the *Ternus temporal ventriloquism* paradigm (Shi, Chen, & Müller, 2010), which we used previously to investigate crossmodal temporal integration. In the standard Ternus temporal ventriloquism paradigm, two auditory beeps are paired with two visual Ternus frames. Visual Ternus displays (see Figure 1) can elicit two distinct percepts of visual apparent motion: *element* or *group* motion, where the type of apparent motion is mainly determined by the visual inter-stimulus interval (ISI_V_) between the two display frames (with other stimulus settings being fixed). Element motion is typically observed with short ISI_V_ (e.g., of 50 ms), and group motion with long ISI_V_ (e.g., of 230 ms) (see Figure 1A, 1B). When two beeps are presented in temporal proximity to, or synchronously with, the two visual frames, the beeps can systematically bias the transition threshold between the two types of visual apparent motion: either towards element motion (if the auditory interval, ISI_A_, is shorter than the visual interval) or towards group motion (if ISI_A_ is longer than the visual interval) (Shi et al., 2010). Similar temporal ventriloquism effects have also been found with other tasks, such as temporal order judgments (for a review, see Chen & Vroomen, 2013). Here, we extended the Ternus temporal ventriloquism paradigm by presenting a whole sequence of beeps prior to the Ternus display frames, in addition to the two beeps paired with Ternus frames (see Figure 1C; recall that previous studies had presented just the latter two beeps) to examine the influence of the temporal averaging of auditory intervals on visual apparent motion.

**Figure 1.**
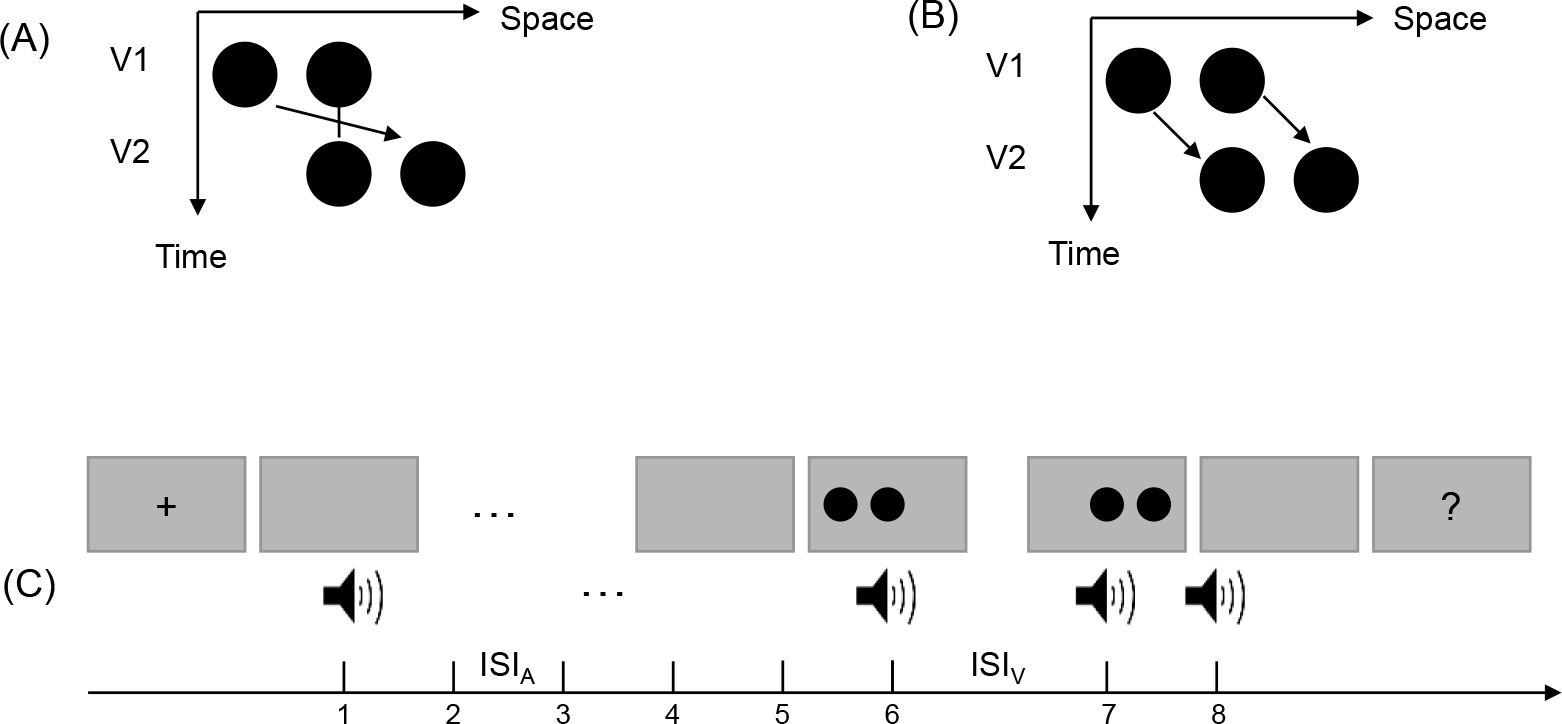
Ternus display and stimulus configurations. Two alternative motion percepts of the Ternus display: (A): ‘element’ motion for short ISIs, with the middle dot perceived as remaining static while the outer dots are perceived to move from one side to the other. (B) ‘group’ motion for long ISIs, with the two dots perceived as moving in tandem. (C) Schematic illustration of the stimulus configurations used in the experiments. The auditory sequence consisted of 8 to 10 beeps. Two of the beeps (the 6th and the 7^th^) were synchronously paired with two visual Ternus frames which were separated by a visual ISI (ISI_V(isual)_) that varied from 50 to 230 ms (for the critical beeps, ISI_V(isual)_ = ISI_A(ditory)_). The other auditory ISIs (ISIA(ditory)) were systematically manipulated such that the mean of the ISI_A_ preceding the visual Ternus display was 50–70 ms shorter than, equal to, or 50–70 ms longer than the transition threshold between the element- and group-motion percepts of the visual Ternus events. The transition threshold was first estimated individually for each observer in a pre-test session. During the experiment, observers were simply asked to indicate the type of visual motion (‘element’ or ‘group’) that they had perceived, while ignoring the beeps.

Experiment 1 was designed, the first instance, to demonstrate an auditory driving effect using this new paradigm. In Experiment 2, we went on to examine whether temporal averaging with irregular auditory sequences would have a similar impact on visual apparent motion. In Experiment 3, we manipulated the variability of the auditory sequence to examine for (and quantify) influences of the variability of the auditory intervals on visual apparent motion. In Experiment 4, we further determined which types of temporal averaging statistics, the arithmetic or the geometric mean of the auditory intervals, influences visual Ternus apparent motion. And Experiment 5 was designed to rule out a potential confound, namely, a ‘recency’ effect – with the last auditory interval dominating the Ternus motion percept – in the crossmodal temporal averaging. Finally, we aimed to identify the computational model that best describes the crossmodal temporal interaction: mandatory full Bayesian integration versus partial integration (Ernst & Banks, 2002; Roach et al., 2006).

## Materials and Methods

### Participants

A total of eighty-four participants (21, 22, 16, 12, 12 in Experiments 1-5; ages ranging from 18–33 years) took part in the main experiments. All observers had normal or corrected-to-normal vision and reported normal hearing. The experiments were performed in compliance with the institutional guidelines set by the Academic Affairs Committee of the Department of Psychology, Peking University (approved protocol of “#Perceptual averaging (2012-03-01)”). All observers provided written informed consent according to the institutional guidelines prior to participating and were paid for their time on a basis of 20 CNY/hour.

The number of participants recruited for Experiments 1 and 2 was based on the effect size in our previous study of the temporal Ternus ventriloquism effect (Shi et al., 2010), where the pairing of auditory beeps with the visual Ternus displays yielded a Cohen’s d greater than 1 for the modulation of the Ternus motion percept. We thus used a conservative effect size of 0.25 and a power of 0.8 for the estimation and recruited more than the estimated sample size (of 15 participants). Given that the effects we aimed to examine turned out quite reliable, we used a standard sample size of 12 participants in Experiments 4 and 5.

### Apparatus and Stimuli

The experiments were conducted in a dimly lit (luminance: 0.09 cd/m^2^) cabin. Visual stimuli were presented in the central region of a 22-inch CRT monitor (FD 225P), with a screen resolution of 1024 × 768 pixels and a refresh rate of 100 Hz. Viewing distance was 57 cm, maintained by using a chin rest.

A visual Ternus display consisted of two stimulus frames, each containing two black discs (l0.24 cd/m^2^; disc diameter and separation between discs: 1.6° and 3° of visual angle, respectively) presented on a gray background (16.1 cd/m^2^). The two frames shared one element location at the center of the monitor, while containing two other elements located at horizontally opposite positions relative to the center (see Figure 1). Each frame was presented for 30 ms; the inter-stimulus interval (ISI_V_) between the two frames was randomly selected from the range of 50–230 ms, with a step size of 30 ms.

Mono sound beeps (1000 Hz, 65 dB, 30 ms) were generated and delivered via an M-Audio card (Delta 1010) to a headset (Philips, SHM1900). To ensure accurate timing of the auditory and visual stimuli, the duration of the visual stimuli and the synchronization of the auditory and visual stimuli were controlled via the monitor’s vertical synchronization pulses. The experimental program was written with Matlab (Mathworks Inc.) and the Psychophysics Toolbox (Brainard, 1997).

### Experimental Design

#### Practice

Prior to the formal experiment, participants were familiarized with visual Ternus displays of either typical element motion (with an ISI_V_ of 50 ms) or typical group motion (ISI_V_ of 260 ms) in a practice block. They were asked to discriminate the two types of apparent motion by pressing the left or the right mouse button, respectively. The mapping between response button and type of motion was counterbalanced across participants. During practice, when a response was made that was inconsistent with the typical motion percept, immediate feedback appeared on the screen showing the typical response (i.e., element or group motion). The practice session continued until the participant reached a conformity of 95%. All participants achieved this criterion within 120 trials, given that the two extreme ISIs used (50 and 260 ms, respectively) gave rise to non-ambiguous percepts of either element motion or group motion.

#### Pre-test

For each participant, the transition threshold between element and group motion was determined in a pre-test session. A trial began with the presentation of a central fixation cross for 300 to 500 ms. After a blank screen of 600 ms, the two Ternus frames were presented synchronized with two auditory tones (i.e., baseline: ISI_V_ = ISI_A_); this was followed by a blank screen of 300 to 500 ms, prior to a screen with a question mark prompting the participant to make a two-forced-choice response indicating the type of perceived motion (element or group motion). The ISI_V_ between the two visual frames was randomly selected from one of the following seven intervals: 50, 80, 110, 140, 170, 200, and 230 ms. There were 40 trials for each level of ISI_V_, counterbalanced with left- and rightward apparent motion. The presentation order of the trials was randomized for each participant. Participants performed a total of 280 trials, divided into 4 blocks of 70 trials each. After completing the pre-test, the psychometric curve was fitted to the proportions of group motion responses across the seven intervals (see *Data Analysis and Modeling*). The transition threshold, that is, the point of subjective equality (PSE) at which the participant was equally likely to report the two motion percepts, was calculated by estimating the ISI at the point on the fitted curve that corresponded to 50% of group motion reports. The just noticeable difference (JND), an indicator of the sensitivity of apparent motion discrimination, was calculated as half of the difference between the lower (25%) and upper (75%) bounds of the thresholds from the psychometric curve.

#### Main Experiments

In the main experiments, the procedure of visual stimulus presentation was the same as in the pre-test session, except that prior to the occurrence of the two Ternus display frames, an auditory sequence consisting a variable number of 6–8 beeps was presented (see below for the details of the onset of the Ternus display frames relative to that of the auditory sequence). As in the pre-test, the onset of the two visual Ternus frames (each presented for 30 ms) was accompanied by a (30-ms) auditory beep (i.e., ISI_V_ = ISI_A_). A trial began with the presentation of a central fixation marker, randomly for 300 to 500 ms. After a 600-ms blank interval, the auditory train and the visual Ternus frames were presented (see Figure 1c), followed sequentially by a blank screen of 300 to 500 ms and a screen with a question mark at the screen center prompting participants to indicate the type of motion they had perceived: element versus group motion (non-speeded response). Participants were instructed to focus on the visual task, ignoring the sounds. After the response, the next trial started following a random inter-trial interval of 500 to 700 ms.

In Experiment 1 (regular sound sequence), the audiovisual Ternus frames was preceded by an auditory sequence of 6–8 beeps with a constant inter-stimulus interval (ISI_A_), manipulated to be 70 ms shorter than, equal to, or 70 ms longer than the transition threshold estimated in the pre-test. The total auditory sequence consisted of 8–10 beeps, including those accompanying the two visual Ternus frames, with the latter being inserted mainly at the 6th–7th positions, and followed by 0–2 beeps (number selected at random), to minimize expectations as to the onset of the visual Ternus frames. Visual Ternus frames were presented on 75% of all trials (504 trials in total). The remaining 25% were catch trials (168 trials) to break up anticipatory processes. All trials were randomized and organized in 12 blocks, each of 56 trials. The ISI_V_ between the two visual Ternus frames was randomly selected from one of the following seven intervals: 50, 80, 110, 140, 170, 200, and 230 ms.

In Experiment 2 (irregular sound sequence), the settings were the same as in Experiment 1, except that the auditory trains were irregular: the ISI_A_ between adjacent beeps in the auditory train (except the ISI_A_ between the beeps accompanying the visual Ternus frames) were varied ±20 ms uniformly and randomly around (i.e., they were either 20 ms shorter or 20 ms longer than) a given mean interval (three levels: 70 ms shorter than, equal to, or 70 ms longer than the individual transition threshold).

Experiment 3 introduced two levels of variability in the auditory-interval sequences with 8–10 beeps: a low coefficient of variance (CV, the standard deviation divided by the mean) of 0.1 and, respectively, a high CV of 0.3. For each CV condition, three arithmetic mean intervals were used: 50 ms shorter than, equal to, or 50 ms longer than the estimated transition threshold. The intervals were randomly generated from a normal distribution with a given mean and CV. The number of the experimental trials was 1008, and the catch trials totaled 336. All trials were randomized and organized in 24 blocks, each block containing 56 trials.

Experiment 4 used three types of auditory sequences, each consisting of 6 intervals: (i) Baseline auditory sequence: three intervals, of 110, 140, and 170 ms, were repeated twice in random order; in this baseline condition, the arithmetic mean (AM = 140 ms) was near-equal to the geometric mean (GM = 138 ms). (ii) AM-deviated sequence: 6 intervals were constructed from ISI_A_ of 70, 140, and 280 ms, which were arranged randomly (AM = 163 ms > GM = 140 ms); (iii) GM-deviated sequence: 6 intervals constructed from ISI_A_ 50, 140, and 230 ms, arranged randomly (GM = 117 ms < AM = 140 ms). The audiovisual Ternus frames were appended at the end of these sequences. The number of experimental trials was 504 (there were no catch trials), which were presented randomized and organized in 12 blocks, each of 42 trials.

To exclude potential confounding by a recency effect, in Experiment 5, we compared two auditory sequences: one with a geometric mean 70 ms shorter than the transition threshold of visual Ternus motion (henceforth referred to as ‘Short’ condition), and the other with a geometric mean 70 ms longer than the transition threshold (‘Long’ condition). Instead of completely randomizing the five auditory intervals (excepting the final synchronous auditory interval with the visual Ternus interval), the last auditory interval before the onset of the Ternus display was fixed at the transition threshold for both sequences. The remaining four intervals were chosen randomly such that the CV of the auditory sequence was in the range between 0.1 and 0.2. This manipulation was expected to minimize the influence of any potential recency effect engendered by the last auditory interval. The audiovisual Ternus frames were appended at the end of these sequences on trials (i.e., 672 out of a total of 784 trials) on which the Ternus display appeared at the end of the sound sequence (the ‘onset’ of the first visual frame was synchronized with 6th beep). The remaining (112) trials were catch trials, with 56 trials each on which the Ternus displays occurred at the beginning of the sound sequence (i.e., the onset of the first visual frame was synchronized with the second beep) or, respectively, at middle temporal locations (i.e., the onset of the first visual frame was synchronized with the 4th beep). These catch trials were introduced to prevent participants from consistently anticipating the visual events to occur at the end of the sound sequence. The total 784 trials were randomized and organized in 14 blocks, each of 56 trials.

### Data analysis and Modeling

We used the R package Quickpsy (Linares & López-Moliner, 2016) to fit psychometric curves with upper and lower asymptotes, which provide better estimates of the thresholds (Wichmann & Hill, 2001). Bayesian modeling was also conducted with R. We first calculated the response proportions for the baseline tests with (audio-) visual Ternus apparent motion and for the formal experiments, as well as fitting the corresponding cumulative Gaussian psychometric functions. Based on the psychometric functions, we could then estimate the discrimination variability of Ternus apparent motion (i.e., *σ_m_*) based on the standard deviation of the cumulative Gaussian function. The parameters of the Bayesian models (see Bayesian modeling section below) were estimated by minimizing the prediction errors using the R *optim* function. Our raw data together with the source code of statistical analyses and Bayesian modeling are available at the github repository: https://github.com/msenselab/temporal_averaging.

## Results

### Experiments 1 and 2: Both regular and irregular auditory intervals alter the visual motion percept

We manipulated the intervals between successive beeps (i.e., the ISI_A_ prior to the Ternus display) to be either regular or irregular, but with their arithmetic mean being either 70 ms shorter, equal to, or 70 ms longer than the transition threshold (measured in the pre-test) between element- and group-motion reports (for both regular and irregular ISI_A_). Auditory sequences with a relatively long mean auditory interval, as compared to a short interval, were found to elicit more reports of group motion, as indicated by the smaller PSEs (Figure 2), for both regular intervals, *F*(2,40)=12.22, *p*<0.001, 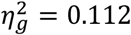, and irregular intervals, *F*(2,42)=8.25, *p*<0.001, 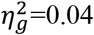. That is, the perceived visual interval (which determines the ensuing motion percept) was assimilated by the average of the preceding auditory intervals, regardless of whether the auditory intervals were regular or irregular. Post-hoc Bonferroni comparison tests revealed that this assimilation effect was mainly driven by the short auditory intervals in both experiments: *ps* were 0.001, 0.00001, and 0.57 for the comparisons −70 vs. 0 ms, −70 vs. 70 ms, and, respectively, 0 vs. 70 ms for the regular intervals; and 0.015, 0.0002, 0.77 for the comparisons of the irregular intervals (Figure 2C and 2D).

**Figure 2.**
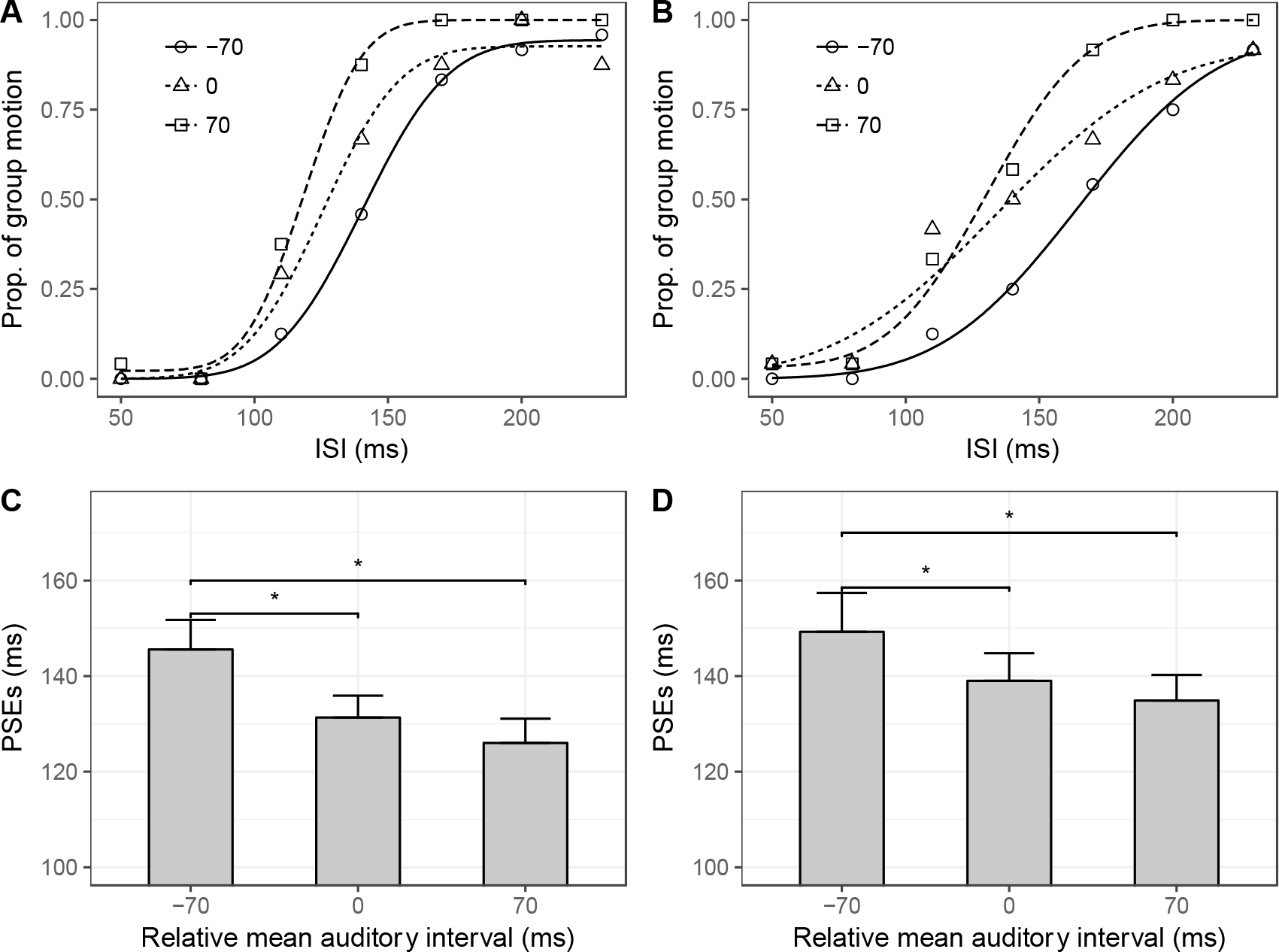
The average means of both regular and irregular auditory sequences influence the visual motion percept. **(A)** Regular auditory-sequence condition: For a typical participant, mean proportions of group-motion responses as a function of the probe visual interval (ISI_v_), and fitted psychometric curves, for auditory sequences with different (arithmetic) mean intervals relative to the individual transition thresholds; the relative-interval labels (−70, 0, and 70) denote the three conditions of the mean auditory interval being 70 ms shorter than, equal to, and 70 ms longer than the pre-test transition threshold, respectively. **(B)** Irregular auditory-sequence condition: for a typical participant, mean proportions of group-motion responses and fitted psychometric curves. **(C)** Mean PSEs as a function of the relative auditory interval for the regular-sequence condition; error bars represent standard errors of the means. **(D)** Mean PSEs as a function of the relative auditory interval for the irregular-sequence condition; error bars represent standard errors of the means.

The fact that a crossmodal assimilation effect was obtained even with irregular auditory sequences suggests that the effect is unlikely due to temporal expectation, or a general effect of auditory entrainment (Jones, Moynihan, MacKenzie, & Puente, 2002; Large & Jones, 1999). In addition, the assimilation effect observed is unlikely due to a recency effect. To examine for such an effect, we split the trials into two categories according to the auditory interval that just preceded the visual Ternus interval: short and long preceding intervals with reference to the auditory mean interval. The length of the immediately preceding interval failed to produce any significant modulation of apparent visual motion, *F*(1, 22) = 2.14, *p* = 0.15. An account in terms of a recency effect was further ruled out by a dedicated control experiment that directly fixed the last auditory interval (see Experiment 5 below).

Furthermore, in the regular condition, the mean JNDs (±SE) for the three ISI_V_ conditions [34.9 (±3.1), 30.5 (±3.4), and 28.4 (±2.9) ms for the ISI_V_ 70 ms shorter, equal to, and, respectively, 70 ms longer relative to the transition threshold] were larger than the JND for the threshold (baseline) condition (18.8 (±1.2) ms; *p*=0.001, *p*=0.002, and *p*=0.033 for the shorter, equal, and longer conditions vs. the ‘threshold’), without differing amongst themselves (all *ps* > 0.1). The same held true for the irregular condition [JNDs of 31.8 (±3.2), *p*=0.001, 30.6 (±2.3), *p*=0.005, and 27.2 (±2.2) ms compared to the baseline 18.6 (±2.1) ms, without differing amongst themselves (all *ps*>0.1). The worsened sensitivities in the three conditions with auditory beep trains suggest that the assimilation effect observed here was not attributable to attentional entrainment, as attentional entrainment would have been expected to enhance the sensitivity.

### Experiment 3: Variability of auditory intervals influences visual Ternus apparent motion

According to quantitative models of multisensory integration (Ernst & Di Luca, 2011; Shi, Church, & Meck, 2013), the strength of the assimilation effect would be determined by the variability of both the auditory intervals and the visual Ternus interval, assuming that information is integrated from all intervals. According to optimal full integration, high variance of the auditory sequence would result in a low auditory weight in audiovisual integration, leading to a weaker assimilation effect compared to low variance. To examine for effects of the variance of the auditory intervals on visual Ternus apparent motion, we directly manipulated the relative standard deviation of the auditory intervals while fixing their arithmetic mean. One key property of time perception is that it is scalar (Church, Meck, & Gibbon, 1994; Gibbon, 1977), that is, the estimation error increases linearly as the time interval increases, approximately following Weber’s law. Given this, we used coefficients of variance (CVs), that is, the ratio of the standard deviation to the mean, to manipulate standardized variability across multiple auditory intervals. Specifically, we compared a low CV (0.1) with a high CV (0.3) condition, with an orthogonal variation of the (arithmetic) mean auditory interval: 50 ms shorter, equal to, or 50 ms longer than the pre-determined transition threshold.

The main effect of mean interval was significant, *F*(2,30)=11.8, *p*<0.001, 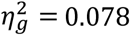, with long intervals leading to more reports of group motion (i.e., lower PSEs: mean PSE of 132±4.6 ms), short intervals to fewer reports of group motion (i.e., higher PSEs: mean PSE of 147±6.7 ms), and equal intervals to an intermediate proportion of group-motion reports (mean PSE of 138±5.3 ms). Post-hoc Bonferroni comparisons revealed this pattern to be similar to that observed in Experiments 1 and 2: significant differences between the short and equal intervals (*p*<0.01) and the short and long intervals (*p*<0.001), but not between the equal and long intervals (p = 0.49). Interestingly, the main effect of CV was significant (though the effect size is small), *F*(1,15) = 5.29, *p*<0.05, 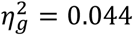, while the interaction between mean interval and CV was not, *F*(2, 30)=0.31, *p*=0.73, 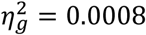 (Figure 3). Further examination for a (potentially confounding) recency effect, adopting the same comparison as for the previous experiments, yielded no evidence that the main effects we obtained are attributable to the length of the auditory interval immediately preceding the visual interval, *F*(1,15) = 0.33, *p* = 0.55.

**Figure 3.**
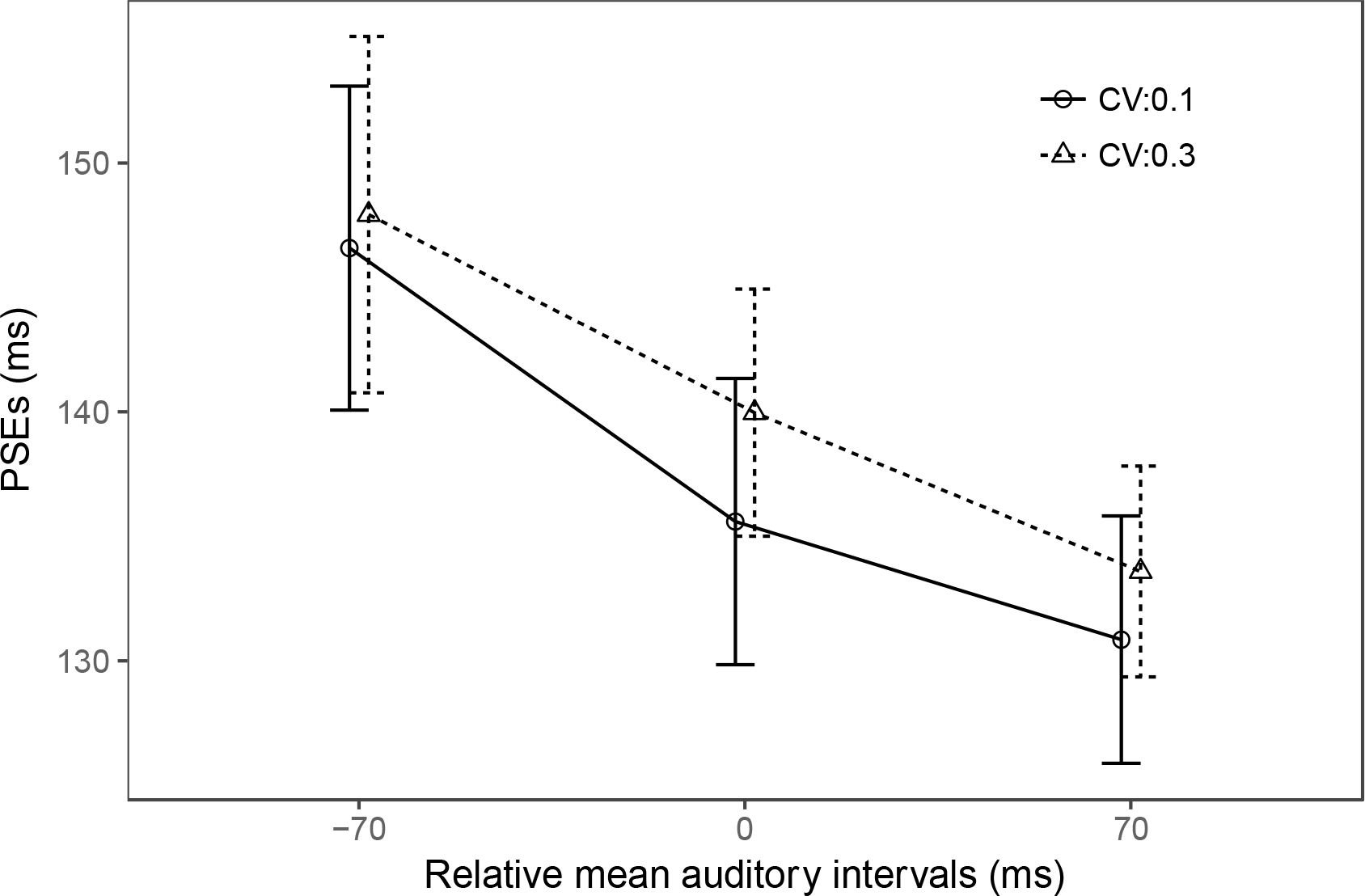
PSEs between element- and group-motion reports for auditory beep trains with a low and a high coefficient of (auditory-interval) variance (CV, 0.1 or 0.3), as a function of the (arithmetic) mean auditory interval (50 ms shorter, equal to, or 50 ms longer than the pre-test transition threshold).

These results are interesting in two respects. First, according to mandatory, full Bayesian integration (see Modeling section below for details), auditory-interval variability should affect the weights of the crossmodal temporal integration (Buus, 1999; Shi et al., 2013), with greater variance lessening the influence of the average auditory interval. Accordingly, the slopes of the fitted lines in Figure 2 would be expected to be flatter under the high compared to the low CV condition, yielding an interaction between mean interval and CV. The fact that this interaction was non-significant suggests that the ensemble mean of the auditory intervals is not fully integrated with the visual interval (we will return to this point in the next, Modeling section). Second, the downward shift of the PSEs in the low, compared to the high, CV condition indicates that the perceived auditory mean interval (that influences the audio-visual integration) is actually not the arithmetic mean (‘AM’) that we manipulated. An alternative account of this shift may derive from the fact the auditory sequences with higher CV have a lower geometric mean (‘GM’) than the sequences with low variance, that is: the perceived ensemble mean is likely geometrically encoded. Experiment 4 was designed to address this (potential) confound by directly comparing the effects of ensemble coding based on the geometric versus the arithmetic mean.

### Experiment 4: Perceptual averaging of auditory intervals assimilates the visual interval towards the geometric, rather than the arithmetic, mean

In Experiment 4, we compared three types of auditory sequence in our audiovisual Ternus apparent motion paradigm: a baseline sequence, an AM-deviated sequence, and a GM-deviated sequence. The PSEs were 136 (±5.46), 148 (±6.17), and 136 (±6.2) ms for the AM-deviated (AriM), the GM-deviated (GeoM), and the baseline conditions, respectively, *F*(2, 22)=8.81, *p*<0.05, 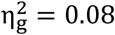 (Figure 4). Bonferroni-corrected comparisons revealed the transition threshold to be significantly larger for the GeoM compared to the baseline condition, *p*<0.01, whereas there was no difference between the AriM and the baseline condition, *p*=1. This pattern indicates that ensemble coding of the auditory interval assimilates the visual interval towards the geometric, rather than the arithmetic, mean.

**Figure 4.**
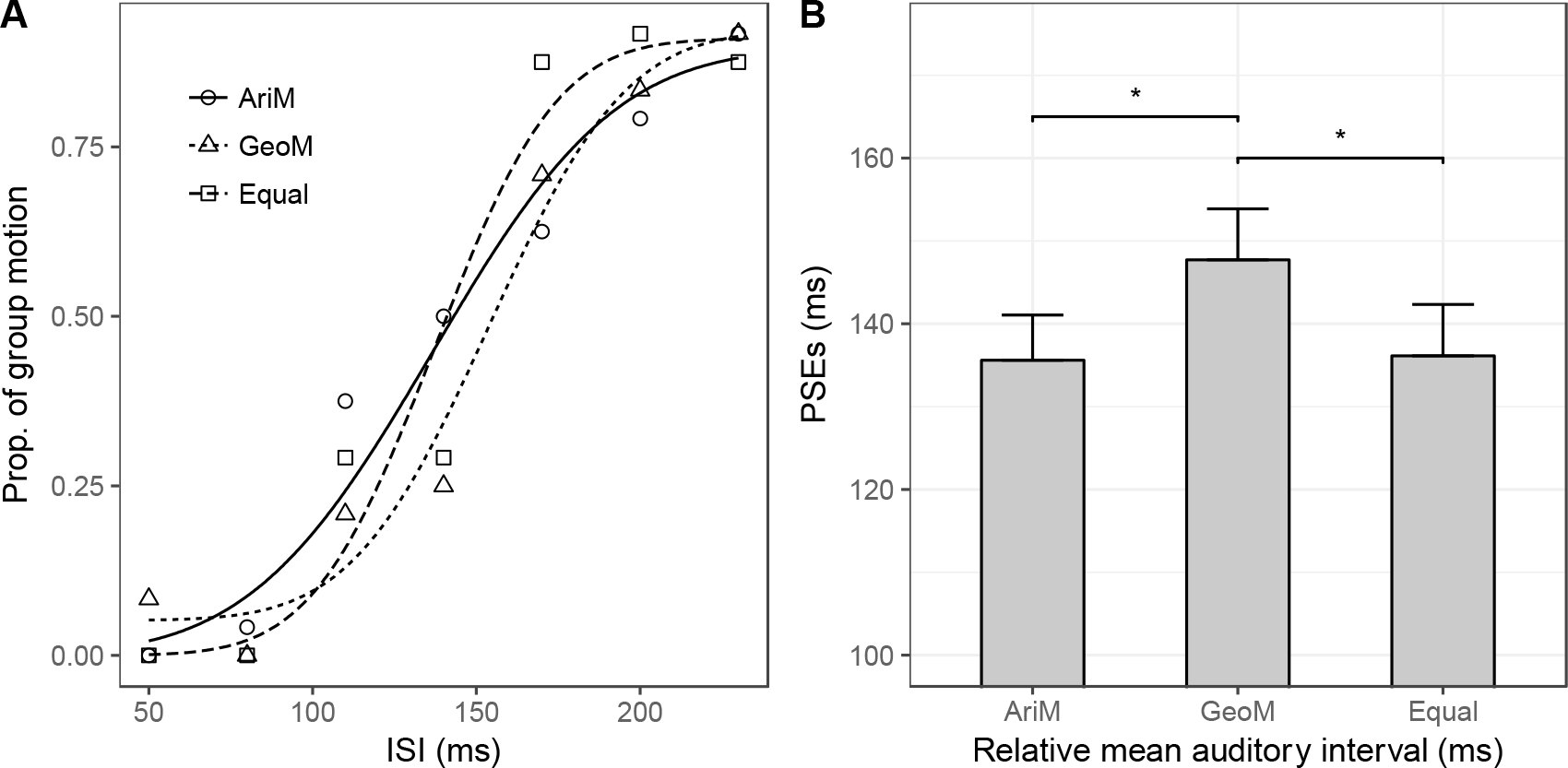
Auditory geometric mean assimilates visual Ternus apparent motion. **(A)** For a typical participant, mean proportions of group-motion responses as a function of the probe visual interval (ISI_v_), and fitted psychometric curves, for the three auditory-sequence conditions: (i) sequence of intervals with larger arithmetic mean (AriM); (ii) sequence of intervals with smaller geometric mean (GeoM); (iii) baseline sequence with equal arithmetic and geometric means (140 ms). **(B)** Mean PSEs (with error bars representing standard errors of the means) for the three auditory-sequence conditions. Compared to the baseline sequence, the GeoM sequence (with the smaller geometric mean) produced a significant shift of the visual transition threshold, whereas the AriM sequence (with the larger arithmetic mean) did not.

### Experiment 5: Auditory sequences with the last interval fixed

In Experiments 1–3, we split the data according to the last interval (i.e., the interval preceding the visual Ternus display) of the auditory sequence into two categories (short vs. long), which failed to reveal any influence of the last interval. In Experiment 5, we formally manipulated the last interval by fixing it at the respective transition threshold for the ‘Short’ and ‘Long’ auditory sequences (i.e., sequences with the smaller and, respectively, larger geometric means). Figure 5 depicts the responses of a typical participant from Experiment 5. The PSEs were 153.1 (±7.3) and, respectively, 137.9 (±9.1) for the ‘Short and ‘Long’ conditions, respectively, *t*(11)=3.640, *p*<0.01. That is, reports of element motion were more dominant in the ‘Short’ than in the ‘Long’ condition, replicating the findings of the previous experiments. In other words, it was the mean auditory interval, rather than the last interval (prior to the Ternus frames), that assimilated visual Ternus apparent motion. Given this, the audiovisual interactions we found here are unlikely attributable to a recency effect.

**Figure 5.**
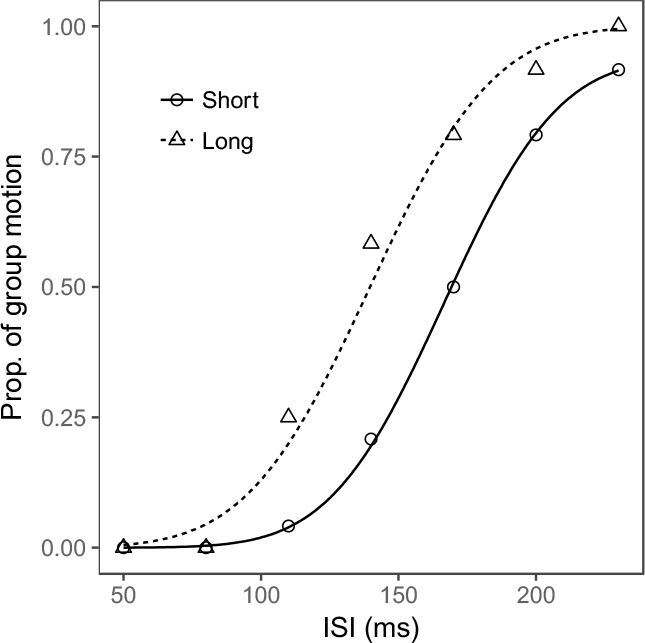
Mean proportions of group-motion responses from a typical participant as a function of the probe visual interval (ISIv), and fitted psychometric curves, for the two geometric mean conditions: the ‘Short’ sequence (with the smaller geometric mean) and the ‘Long’ sequence (with the larger geometric mean).

## Bayesian modeling

To account for the above findings, we implemented, and compared two variants of Bayesian integration models: mandatory full Bayesian integration and partial Bayesian integration. If the ensemble-coded auditory-interval mean (*A*) and the audiovisual Ternus display interval (*M*) are fully integrated according to the maximum likelihood estimation (MLE) principle (Ernst & Banks, 2002), and both are normally distributed (e.g., fluctuating due to internal Gaussian noise) – that is: *A* ~ *N*(*I_a_, σ_a_*), *M* ~ *N* (*I_m_, σ_m_*) – the expected optimally integrated audio-visual interval, which yields minimum variability, can be predicted as follows:

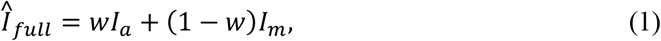

where 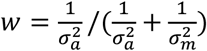 is the weight of the averaged auditory interval, which is proportional to its reliability. Note that full optimal integration is typically observed when the two ‘cues’ are close to each other, but it breaks down when their discrepancy becomes too large (Kording et al., 2007; Parise, Spence, & Ernst, 2012; Roach et al., 2006). In our study, the Ternus interval and the mean auditory interval could differ substantially on some trials (e.g., visual interval of 50 ms paired with mean auditory interval of 210 ms). Given this, a more appropriate model would need to take a ‘discrepancy’ prior and the causal structure (Kording et al., 2007) of audio-visual temporal integration into consideration. Thus, similar to (Roach et al., 2006), here we assume that the probability of full integration *P_am_* depends on the discrepancy between the mean auditory and Ternus intervals:

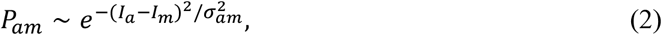

where 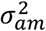 is the variance of the sensory measures of the discrepancy between the ensemble mean of the auditory intervals and the visual interval. *P_am_* will vary from trial to trial, depending on the discrepancy between the mean auditory interval and the visual interval. Thus, a more general, partial integration model would predict:

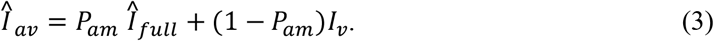

Combined with equation (1), equation (3) can be simplified as follows:

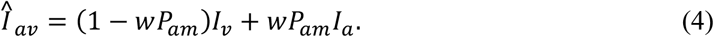

To compare the full-integration and partial-integration models, we took into account the data from those of our experiments that manipulated the auditory-interval regularity and variability (Experiments 1–3; we excluded Experiments 4 and 5, as these did not include a baseline task of Ternus apparent-motion perception; see Methods section). Given that the baseline task provided an estimate of *σ_m_*, there is one parameter – *σ_a_* – for the full-integration model and two parameters – *σ_a_* and *σ_am_* – for the partial-integration model, which require parameter fitting. This was carried out using the optimization algorithm L-BFGS in R (see our source code at https://github.com/msenselab/temporal_averaging). We assessed the goodness of the resulting fits by means of coefficients of determination (*R*^2^) and Bayesian information criteria (*BIC*). The *BIC* and *R*^2^ scores are presented in Table 1. As can be seen, the BIC differences between the partial- and full-integration models are large for all experiments, clearly favoring the partial-integration model (Kass & Raftery, 1995). The *R*^2^ values also confirms this.

**Table 1.** Model comparison using BIC and ***R*^2^** for the partial- and full-integration model.

| Experiments | Partial Integration | | Full Integration | | $\Delta BIC$ |
| --- | --- | --- | --- | --- | --- |
| | $BIC$ | $R^2$ | $BIC$ | $R^2$ | |
| **Irregular** | -1859 | 0.86 | -1392 | 0.63 | **467** |
| **Regular** | -1932 | 0.91 | -1772 | 0.88 | **160** |
| **Variance** | -2894 | 0.91 | -2878 | 0.91 | **16** |

*Note. The differential BICs scores revealed the partial-integration model to outperform the full-integration model across all experiments (very strong evidence in all experiments: $\Delta BIC > 10$ ).*

To visualize how well the partial-integration model predicts behavioral performance, we calculated the predicted mean responses based on the partial-integration model for individual visual ISIs across all experimental conditions. Figure 6 illustrates the predictions, indicated by curves, together with the observed mean responses, indicated by shape points. As can be seen, the predicted mean responses are within one standard error of the observed mean responses (Figure 6).

**Figure 6.**
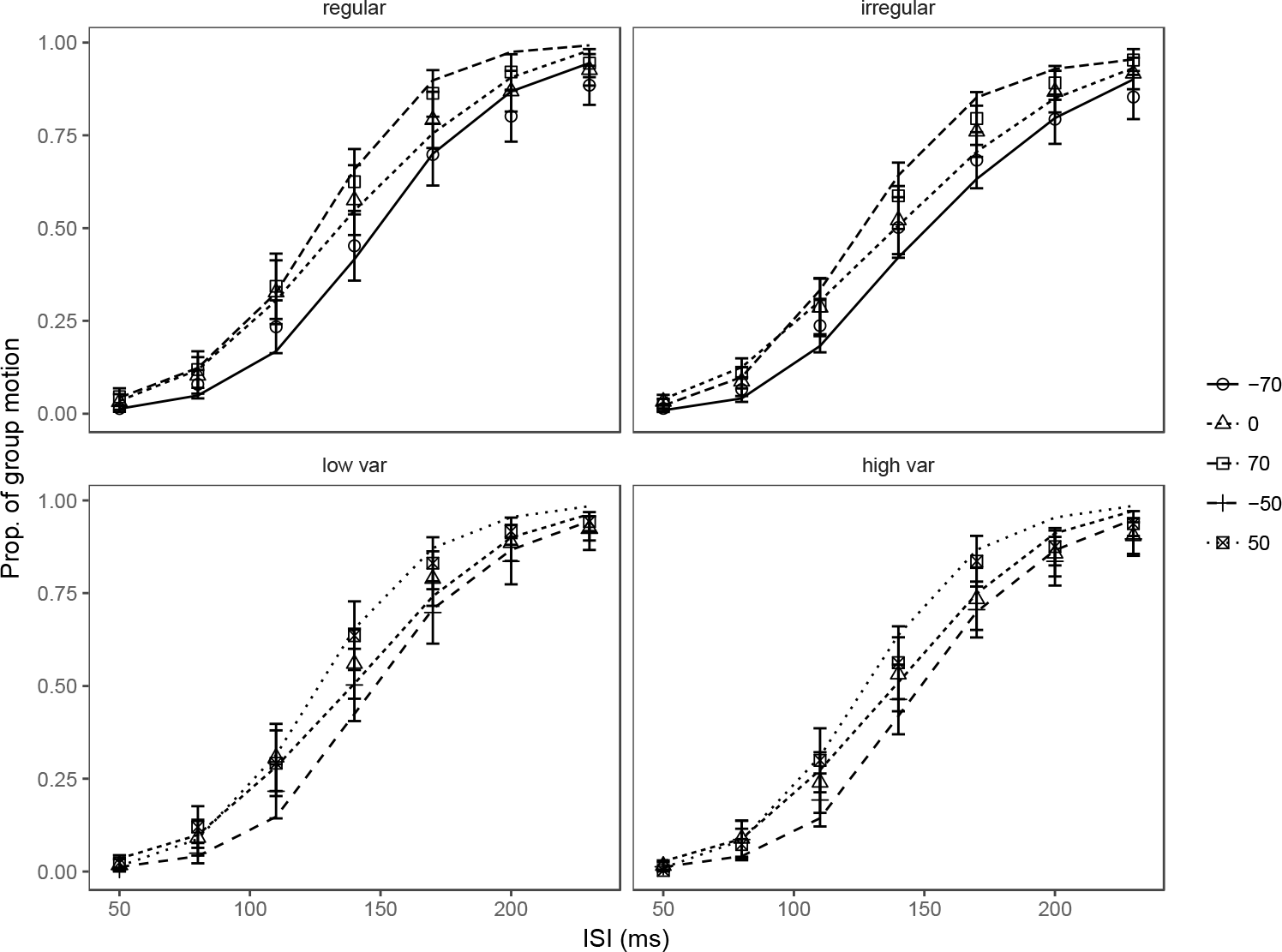
Mean behavioral responses (proportion of group-motion reports, indicated by shape points) and responses predicted by the partial-integration model (indicated by curves) as a function of the ISI_V_ of the Ternus display, separately for auditory sequences with different (arithmetic) mean intervals relative to the individual transition thresholds. The relative-interval labels (−70, −50, 0, 50, and 70 [ms]) denote the magnitude of the difference between the mean auditory interval and the transition threshold. Error bars denote standard errors of means (±SEM).

The key difference between the full- and partial-integration models is that the latter takes the probability of crossmodal integration into account; accordingly, the weight of the auditory ensemble intervals (i.e., *wP_am_*) depends on the difference between the ensemble mean of the auditory intervals and the visual interval. This can be seen in Figure 7, which illustrates the dynamic changes of the auditory weights across the various audio-visual interval discrepancy conditions. All three experiments exhibit a similar pattern: weights are at their peak when the visual interval and the auditory mean intervals are close to each other. For example, the peaks for the relative intervals of 0 ms (i.e., the auditory mean intervals were set to the individual visual thresholds) are around 140 ms, close to the mean visual transition threshold (134.6 ms for regular and 135.3 ms for irregular sequences, and 139.0 ms for low and 144.8 ms for high variance). For relative intervals of 70 ms, the peaks are shifted rightwards; and for relative intervals of −70 ms, they are shifted leftwards.

**Figure 7.**
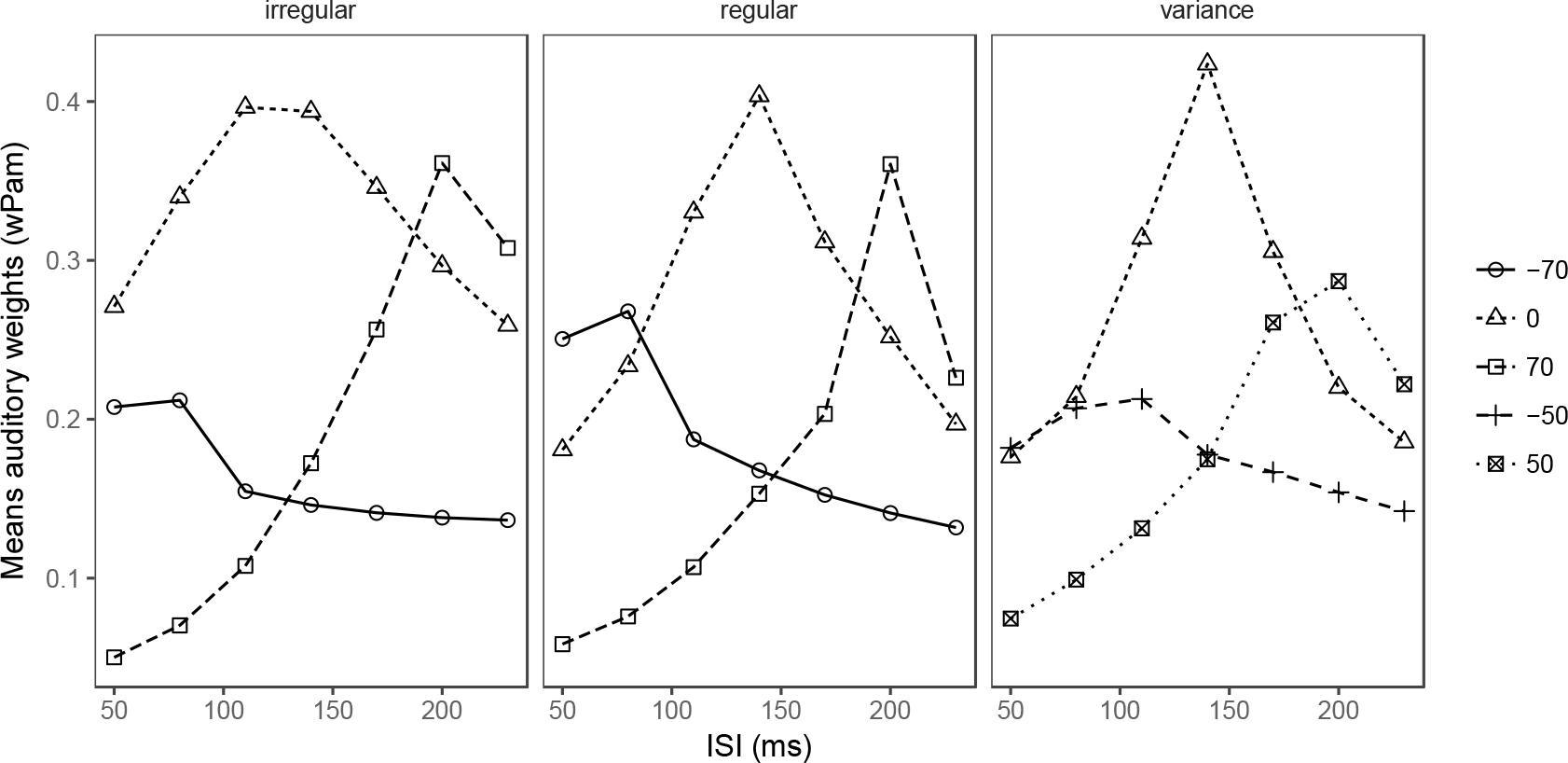
Predicted weights (i.e., *wP_am_*, based on the partial-integration model) of the auditory ensemble intervals as a function of the ISI_V_ of the Ternus display, separately for auditory sequences with different (arithmetic) mean intervals relative to the individual transition thresholds. The relative-interval labels (−70, −50, 0, 50, and 70 ms) denote the magnitude of the difference between the mean auditory interval and the transition threshold.

Based on the responses predicted by the partial-integration model, we further calculated the predicted PSEs. Figure 8 shows a linear relation between the observed and predicted PSEs for all experiments. Linear regression revealed a significant linear correlation, with a slope of 0.978 and an adjusted *R^2^* = 0.983. The full-integration model, by contrast, produced flat psychometric curves for 6% of the individual conditions in Experiments 1 and 2 (due to the weight of the mean auditory interval approaching 1), which yielded unreliable estimates of the corresponding PSEs. This led to lower predictive power compared to the partial-integration model, as evidenced by the BIC and R^2^ scores (Table 1). Thus, taken together, the partial-integration model can well explain the behavioral data that we observed.

**Figure 8.**
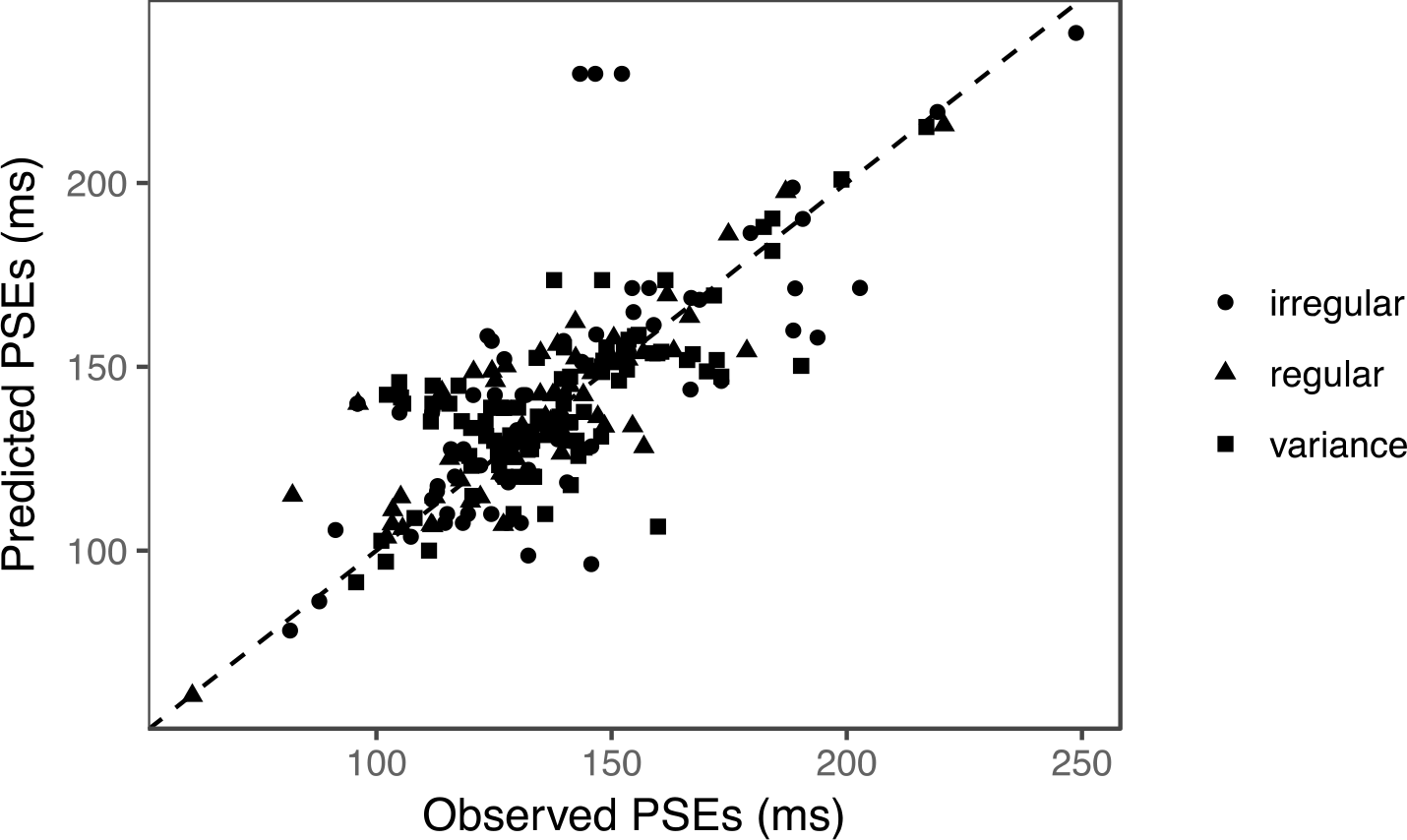
Predicted PSEs versus observed PSEs for all experiments. Each dot represents the PSE of one particular observer in a given experimental condition. Shape points represent the four auditory-sequence manipulations. Linear regression revealed a significant high correlation (*R*^2^ = 0.983) and a slope of 1.008.

## General Discussion

Using an audiovisual Ternus apparent motion paradigm, we conducted five experiments on audiovisual temporal integration with regular and irregular auditory sequences presented prior to the (audio-) visual Ternus display. We found that perceptual averaging of both regular (Experiment 1) and irregular auditory sequences (Experiments 2 and 3) greatly influenced the timing of the subsequent visual interval, as expressed in systematic changes of the transition threshold in visual Ternus apparent motion: longer mean auditory intervals elicited more reports of group motion, whereas shorter mean intervals gave rise to dominant element motion. In Experiment 4, we further found that the geometric mean of the auditory intervals can explain the audiovisual interaction better than the arithmetic mean. Further (post-hoc) analyses and a purpose-designed experiment (Experiment 5) effectively ruled out an explanation of these findings in terms of a recency effect, that is, a dominant influence of the last interval prior to the Ternus frames. Using a Bayesian integration approach, we showed that the behavioral responses are best predicted by partial-cue integration, rather than by full integration. Thus, our results reveal the processing – in particular, the temporal averaging – of a train of beeps that forms the background context of the visual task to play a critical role in crossmodal temporal integration, even when participants are asked to ignore the auditory stimuli.

### Perceptual averaging and crossmodal temporal rate interaction

Extracting key statistical information from sets of objects or events in our environment would provide us with a perceptual strategy to cope with limitations in attentional and working memory capacity (Allik, Toom, Raidvee, Averin, & Kreegipuu, 2014; Chetverikov, Campana, & Kristjansson, 2016) – given that we can have conscious access to only very few items from the total amount of information received by our senses at any one time (e.g., Bundesen, Habekost, & Kyllingsbaek, 2005; Cohen, Dennett, & Kanwisher, 2016; Cowan, 2001; Marois & Ivanoff, 2005). In this situation, perceptual averaging would endow us with an efficient and, in evolutionary terms, competitive solution to overcome bandwidth limitations (McClelland & Bayne, 2016), thus constituting one of the underlying computational principles for selecting appropriate actions to achieve our current behavioral goals. Clearly, timing is fundamental for dynamic perception, and therefore unlikely to be an exception with regard to perceptual averaging (Hardy & Buonomano, 2016; McDermott & Simoncelli, 2011). For instance, when listening to a piece of music, we can immediately tell the average tempo, even though the individual ‘notes’ may not be well remembered. And when watching a field of runners in a competition, we immediately know whether it is a slow or a fast race overall.

Research on the audiovisual interaction in (crossmodal) event timing has shown auditory rate to have a pronounced influence on visual rate perception (Recanzone, 2003, 2009; Roach et al., 2006; Shipley, 1964). The visual temporal rate is often assimilated to the auditory rate, owing to the higher temporal resolution of audition compared to vision. Of note, however, the extant studies have used only regular temporal sequences, thus leaving it an open question whether the mechanism underlying the assimilation effect is perceptual averaging, temporal entrainment, or a recency effect from the latest auditory interval. On this background, the present study examined how irregular auditory sequences influence visual interval timing – measured in terms of the transition threshold of Ternus apparent motion – and showed that it is the temporal averaging of the auditory sequence (regardless of its regularity) that exerted a great influence on the visual interval.

### Temporal averaging and geometric encoding

The present results indicate that the geometric mean well encapsulates the summary statistics of the temporal structure hidden in a complex multisensory stream (Hanson, Heron, & Whitaker, 2008; Heron, Roach, Hanson, McGraw, & Whitaker, 2012). Previous work on numerosity had already suggested that the mental scales underlying the representation of visual numerosity and temporal magnitudes are best characterized as being non-linear, as opposed to linear, in nature (Dehaene, 2003; Dehaene et al., 2008; Nieder & Miller, 2003, 2004; Rips, 2013). For example, adults from the Mundurucu, an Amazonian indigenous tribe with a limited number lexicon, map numerical quantities onto space in a logarithmic fashion (Dehaene et al., 2008; but see Cicchini, Arrighi, Cecchetti, Giusti, & Burr, 2012). A seminal study by Allan and Gibbon also showed that temporal bisection coincided with the geometric mean of the two reference durations (Allan & Gibbon, 1991). Our findings reveal that extraction of the geometric mean also underlies temporal averaging – and this might well be a principle shared by a broad range of mechanisms coding ‘magnitude’ in perception (Walsh, 2003).

### Partial integration in crossmodal temporal processing

Research on multisensory integration has shown that the ‘proximity’ and ‘similarity’ of the spatiotemporal structure of multisensory signals – technically, their cross-correlation in time (and space) – is critical for inferring an underlying common source to both signal streams (Parise & Ernst, 2016; Parise et al., 2012). Accordingly, highly correlated audiovisual events are likely perceived as arising from a single, multisensory source. Roach and colleagues (2006) quantified this for audiovisual rate perception by introducing a disparity prior, that is, their model assumes that the strength of crossmodal temporal integration is dependent on the disparity between the auditory and visual temporal rates.

In the present study, by comparing two variants of Bayesian integration models, full and partial integration, our findings also quantitatively elucidate the way in which geometric averaging of the preceding, task-irrelevant auditory intervals assimilates the subsequent, perceived visual interval between the Ternus display frames. The modeling results indicate that the ensemble mean of the auditory intervals only *partially* integrates with the visual interval, dependent on the time disparity between the two: when the mean of the auditory intervals is close to the visual interval, they are optimally integrated according to the MLE principle; in contrast, if the ensemble mean deviates grossly from the visual interval, partial integration, based on the crossmodal disparity, provides a superior account of the behavioral data to mandatory, full integration. However, in contrast to full integration, partial integration requires participants to take both the mean statistics and the crossmodal disparity into account. This is consistent with a large body of literature on temporal contextual modulation, within the broader framework of Bayesian optimization (Jazayeri & Shadlen, 2010; Roach, McGraw, Whitaker, & Heron, 2017; Shi et al., 2013), where prior information (e.g., history information or a discrepancy prior) is incorporated in multisensory integration.

### Perceptual averaging and temporal entrainment

One important question to be considered is whether the assimilation effect induced by perceptual averaging can be distinguished, at root, from attentional entrainment. In the typical auditory entrainment paradigm, the rhythm itself is irrelevant with respect to the visual target events that are to be detected (or discriminated), though temporal expectations induced by the rhythm influence attentional selection of the target (Lakatos, Karmos, Mehta, Ulbert, & Schroeder, 2008). Rhythmically (i.e., with temporal attention) anticipated target events are detected or discriminated more rapidly than early or late events that are out of phase with the peaks of the attentional modulation induced by the entrainment (Ronconi & Melcher, 2017). Irregular rhythms, by contrast, have been shown to disrupt temporal attention, as evidenced by reduced benefits for responding to the target events (Miller, Carlson, & McAuley, 2013). Importantly, in present study, both regular and irregular auditory sequences did reduce (rather than enhance) the sensitivity of discriminating Ternus apparent (i.e., element vs. group) motion, as evidenced by the increased JNDs. In contrast, the averaged temporal intervals, whether these formed a regular or irregular series, were automatically integrated with the subsequent visual interval, as expressed in the systematic biasing of the reported visual motion percepts. This ‘dissociation’ implies that the assimilation effects demonstrated here reflect a genuine, automatic perceptual averaging mechanism that operates independently of attentional entrainment processes.

### Irrelevant context in multisensory perceptual averaging

One might ask why the brain would at all take into account entirely task-irrelevant contexts – such as, in the present study, the (mean of the) intervals of an irrelevant auditory sequence – in multisensory integration. As revealed by our experiments, the discrimination sensitivity for visual apparent motion became actually worse and the motion percept became biased by including the irrelevant auditory sequence. Note however that, in the real world, there are normally strong associations and correlations in the multisensory inputs – so that drawing on this additional information often increases the reliability of perceptual estimates. For example, the rhythmic sound pattern produced by a train moving along the track would help us improve our estimation of the train’s speed, given that the tempo of the track sound is linearly correlated with the speed of the train. Indeed, convergent evidence suggests that multisensory integration can reduce the uncertainty of the final estimates in many situations (Ernst & Banks, 2002; Ernst & Di Luca, 2011). However, integrating multiple sources of information that deviates from the currently relevant information may engender unwanted biases. Such contextual modulations have been reported in various forms. For example, when performing a series of time estimations, observers’ judgment of a given interval is biased toward the intervals that they just experienced (Jazayeri & Shadlen, 2010) – which is known as a *central-tendency effect* (Petzschner, Glasauer, & Stephan, 2015; Shi & Burr, 2016; Shi et al., 2013). A similar contextual modulation is also at work in the so-called *time-shrinking illusion*, in which the percept of the last auditory interval is assimilated by the preceding intervals (Nakajima, ten Hoopen, Hilkhuysen, & Sasaki, 1992; Nakajima et al., 2004), as well as in audiovisual interval judgments when auditory and visual intervals are presented sequentially (Burr et al., 2013). The present study demonstrated that such an audiovisual integration still occurs even when participants are explicitly told to ignore the (task-irrelevant) auditory sequence, suggesting that processes of top-down control cannot fully shield visual motion perception from audiovisual temporal integration.

## Conclusion

It has long been known that auditory flutter drives visual flicker (Shipley, 1964) – a typical phenomenon of audiovisual temporal interaction with regular auditory sequences. Here, in five experiments, we demonstrated that irregular auditory sequences also capture temporal processing of subsequently presented visual (target) events, measured in terms of the biasing of Ternus apparent motion. Importantly, it is the geometric averaging of the auditory intervals that assimilates the visual interval between the two visual Ternus display frames, thereby influencing decisions on perceived visual motion. Further work is required to examine whether the principles of geometric averaging and partial crossmodal integration demonstrated here (for an audiovisual dynamic perception scenario) generalize to other perceptual mechanisms underlying magnitude estimation in multisensory integration.

### Context of the Research

Perceptual averaging of sensory properties, such as the mean number, size, and spatial layout of objects in a scene, has been documented extensively in the visuospatial domain. It allows us to capture our environment at a glance, in summary terms – overcoming attentional and working-memory capacity limitations. This phenomenon prompted us to ask whether and, if so, how processes of perceptual averaging may also be applied in the temporal domain, specifically in (crossmodal) scenarios involving multiple interacting sensory systems. Thus, we designed a paradigm combining a task-irrelevant temporal sequence of auditory events with task-relevant Ternus apparent motion – a phenomenon where we see two aligned dots either move together (e.g., to the left or right) or only one dot ‘jumping’ across the other (apparently stationary) dot. What we see (group vs. element motion) is critically influenced by the temporal interval between the two Ternus display frames. What we found is that the irrelevant auditory sequence preceding the visual Ternus display alters the visual interval, thus biasing observers to see either more group motion or more element motion, depending the geometric mean of the preceding auditory intervals. This interaction depends on the discrepancy between the (mean) auditory and the visual interval: if the discrepancy becomes too large, no interaction occurs. Conceptually, the finding of temporal averaging over a sequence of auditory intervals and its subsequent influence on the visual interval makes a connection to the psychophysically well-established *central-tendency effect*, in which the prior sampled distribution – here: of the auditory intervals – ‘assimilates’ the estimate – here: the visual interval. Although we have provided a formal (partial Bayesian integration) description of this crossmodal assimilation effect, further, purpose-designed research is required to provide a complete picture of underlying, interacting neural mechanisms.

## Notes

Conflict of Interest: None

